# Tuning Diversity Improves Discrimination and Detection Performance under Metabolic Constraints

**DOI:** 10.64898/2026.06.29.735317

**Authors:** Dario L. Ringach

**Affiliations:** Departments of Neurobiology & Psychology, David Geffen School of Medicine, University of California, Los Angeles, Los Angeles, CA 90095, USA

**Keywords:** population coding, tuning diversity, orientation tuning, discrimination, detection, metabolic constraints, information tuning curve, efficient coding

## Abstract

Cortical populations exhibit a wide range of tuning properties, raising the question of whether such variability is a feature or a bug of cortical function. Prior work has shown that tuning diversity can improve population codes by mitigating the effects of correlated noise and increasing the discrimination and identification capacity of geometric representations. Motivated by these findings, we study a model in which a heterogeneous family of tuning curves, coding for a circular variable, is replicated at equally spaced preferred angles. We show that this heterogeneous population achieves better discrimination and detection than an equally sized homogeneous population constructed from shifted copies of the family’s mean tuning curve, while using the same spike budget. Thus, homogeneous tuning is unstable under perturbations that preserve the mean tuning curve, because such perturbations leave metabolic cost unchanged while improving coding performance. We propose that such instability creates evolutionary pressure toward heterogeneity of tuning, making its prevalence a consequence of a process that optimizes coding performance under metabolic constraints.

**Significance Statement:** The tuning curves of cortical neurons vary substantially, raising the question of whether this diversity has functional significance or merely reflects biological noise. We show that perturbing an initially homogeneous population while preserving its mean tuning curve produces a heterogeneous population with better discrimination and detection performance at the same metabolic cost. Thus, tuning diversity may emerge as a natural consequence of selection for efficient coding under metabolic constraints.

## Introduction

Neurons in primary visual cortex exhibit substantial diversity in receptive-field organization, orientation and direction selectivity, spatial-frequency tuning, and contrast-response properties [1–7]. This functional diversity is paralleled by a continuous distribution of transcriptomic signatures in visual cortical neurons [8]. Whether such diversity should be understood as nuisance variability or as a functional feature of population coding remains an important theoretical question. It has been conjectured, for example, that a continuous distribution of cell types between extreme archetypes may serve as a mechanism for division of labor among cells that collectively perform multiple tasks under trade-offs [9]. Prior results show that diversity can indeed be beneficial to cortical coding of orientation: it can prevent information from saturating in the presence of structured correlations [10], alter the effects of noise covariance in ways missed by homogeneous models [11, 12], and improve the discrimination and identification capacity of geometric representations [13].

Extending this line of work, here we study a simplified model of heterogeneity in a neural population coding for a circular variable. A finite family of tuning curves is replicated at equally spaced preferred orientations (with grid spacing Δ), so that coverage of the circle is complete and identical for each tuning-curve type. This construction removes sampling asymmetries and isolates the effect of tuning diversity itself, and makes the model mathematically tractable. We compare this heterogeneous replicated population with a homogeneous one consisting of the same number of neurons whose tuning curve is the mean in the heterogeneous population. Thus, metabolic cost, measured by the expected *L*^1^ norm (number of spikes), remains fixed. For orientation coding, a useful performance measure is the information tuning curve: *d*^′2^(*r*Δ) of two population responses as a function of angular separation (here restricted to multiples of the grid spacing) [14].

The first set of results concerns *discrimination* between two stimuli. Under independent, isotropic Gaussian noise, raw *d*^′2^ is proportional to the squared Euclidean distance between population response vectors. Here, we show that perturbations in the shape of tuning curves that preserve the mean tuning curve of a homogeneous population improve discrimination at *all* angular separations. The same result can be obtained for independent Poisson counts using a finite-separation measure based on Chernoff information between the two population count distributions [14]. The second set of results concerns *detection* of a stimulus from baseline activity. We assume that, in the absence of a stimulus, every neuron fires at a common baseline rate *r*_0_. For both Gaussian and Poisson noise, we show that detection in the heterogeneous population is better than in the homogeneous one.

These discrimination and detection results are the main focus of the manuscript. The main message is that, in a simple model of population heterogeneity, homogeneous tuning is unstable with respect to perturbations that leave the mean tuning curve invariant: such perturbations preserve the metabolic cost, measured by total spike budget, while improving performance. This instability creates evolutionary pressure toward heterogeneity of tuning, making its prevalence a consequence of a process that optimizes coding performance under metabolic constraints.

## Results

### Discrimination under independent Gaussian noise

We first consider discrimination between two stimulus orientations under independent Gaussian noise. Let *f*_*q*_[*n*], *q* = 1, …, *Q*, denote a finite family of nonnegative tuning curves sampled on an equally spaced circular grid *n* = 0, …, *K* − 1. Each tuning-curve type is replicated at every preferred orientation. Thus, for a stimulus at grid position *j*, the heterogeneous population response contains all circular shifts of the curves *f*_*q*_.

For independent Gaussian noise with common variance *σ*^2^, squared discriminability between two response vectors is proportional to their squared Euclidean distance. For orientations separated by *r* grid steps, the raw Gaussian information curve is

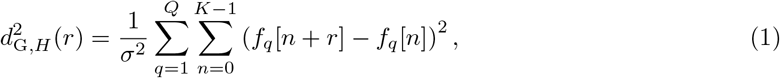

where indices are taken modulo *K*. Because the population contains all preferred orientations, this quantity is independent of the reference orientation *j*.

Now define the arithmetic mean tuning curve

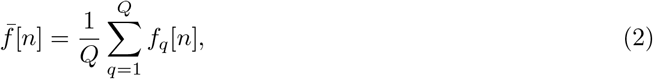

and compare the heterogeneous population with a homogeneous reference population consisting of *Q* copies of 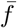, again replicated at all preferred orientations. The corresponding information curve is

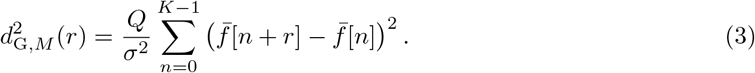

Write each tuning curve as the mean plus a deviation,

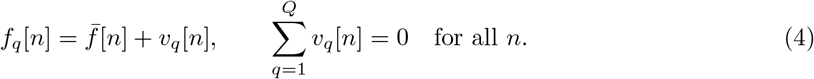

Then

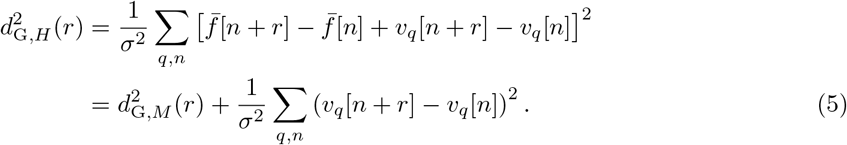

The cross-term vanishes because the deviations sum to zero at each orientation. Therefore

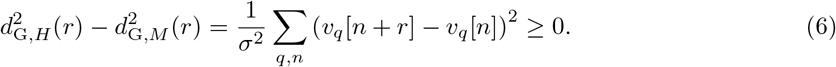

Thus the heterogeneous population cannot have lower raw Gaussian discrimination performance than *Q* copies of its mean tuning curve at any angular separation. Since the tuning curves are non-negative, the total *L*^1^ metabolic cost is the same:

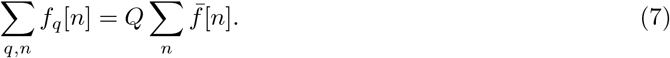

Thus, under an *L*^1^ spike-budget constraint, the heterogeneous population weakly dominates the mean-copy homogeneous population.

The equality condition in Eq. 6 holds if and only if, for any given *r*,

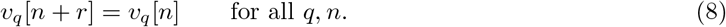

That is, the deviations must be invisible to the shift being tested. Only DC perturbations are neutral for the full information curve. Thus, any non-constant mean-preserving perturbation striclty improves performance for at least some angular separations.

### Discrimination under independent Poisson noise

For independent Poisson responses, finite-separation discrimination should be defined from likelihoods rather than from Fisher information alone. Let two stimuli produce independent Poisson count distributions with rate vectors *λ* and *μ*. The Chernoff information is

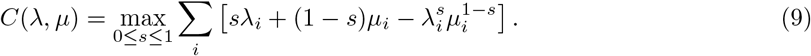

The factor 8*C* is a natural Gaussian-calibrated *d*^′2^, since the Chernoff information of two equal-covariance Gaussian distributions is one eighth of their squared Mahalanobis distance.

For the replicated heterogeneous family, define

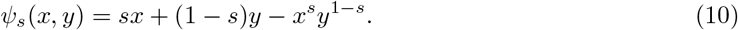

Then

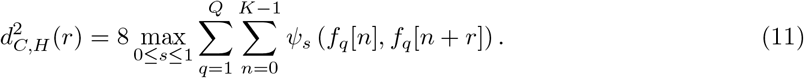

We assume the rates are striclty positive. For fixed *s* ∈ (0, 1), the function *x*^*s*^*y*^1−*s*^ is concave and homogeneous of degree one, so *ψ*_*s*_ is convex. Jensen’s inequality gives

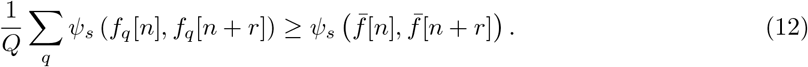

Summing over *n* and maximizing over *s* yields

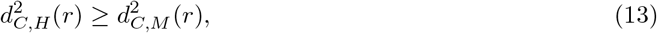

where *M* denotes *Q* replicated copies of the mean tuning curve. The total expected spike count is the same in the two populations,

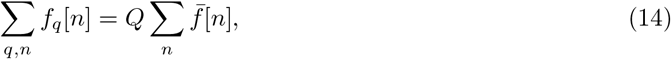

so under *L*^1^ firing-rate cost the heterogeneous family again weakly dominates the mean-copy population.

The equality condition differs from the Gaussian case. For fixed *s* ∈ (0, 1), equality in Jensen holds at a particular *n* and *r* if and only if the points

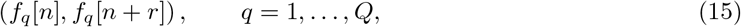

lie on a common ray from the origin. Equivalently, the ratio

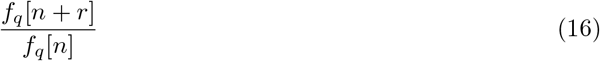

must be independent of *q*. If the mean-copy Chernoff information at separation *r* is nonzero, its maximizing *s* is interior, so equality of the maximized Chernoff *d*^′2^ at that separation requires this proportional-pair condition for every *n*. For equality of the whole information curve, a simple positive-rate case is that all tuning curves are multiplicative copies of one common shape,

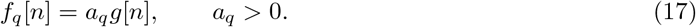

Thus pure gain diversity is neutral for the Poisson Chernoff comparison with *Q* copies of the mean. Shape diversity, in contrast, gives a strict improvement whenever it violates the proportional-pair condition at the relevant separation.

### Stimulus detection under Gaussian noise

Now consider detection rather than discrimination. In the no-stimulus condition, we assume every neuron has baseline rate *r*_0_. When a stimulus is present at orientation *j*, the heterogeneous population response is the complete set of shifted rates *f*_*q*_[*j* − *c*]. Since changing *j* only permutes the components of the response vector, detection performance is the same for every orientation. Thus the average *d*^′^ over orientations is simply the common value of *d*^′^.

In the independent Gaussian case with unit variance, the squared detection index for the heterogeneous population is

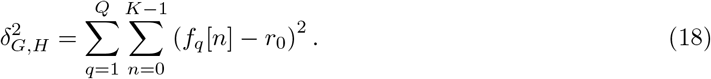

For *Q* copies of the mean tuning curve,

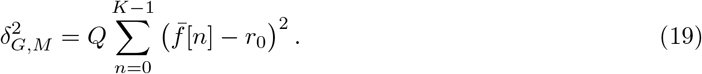

Using 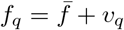 and ∑_*q*_ *v*_*q*_ = 0 pointwise in *n*,

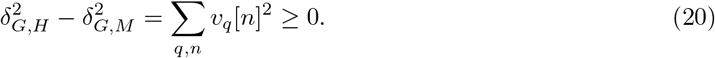

Therefore heterogeneity improves Gaussian detection relative to the mean-copy population. Equality holds if and only if *v*_*q*_[*n*] = 0 for every *q* and *n*, meaning the tuning curves are identical on the sampled grid. For a genuinely heterogeneous population, we have strict improvement in detection.

### Stimulus detection under Poisson noise

For Poisson detection, the Chernoff information between a stimulus rate *x* and baseline rate *r*_0_ uses

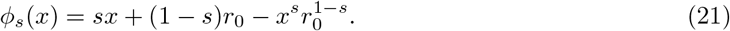

The heterogeneous and mean-copy Chernoff exponents are

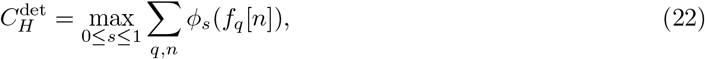

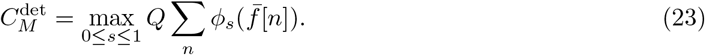

For *r*_0_ *>* 0 and *s* ∈ (0, 1),

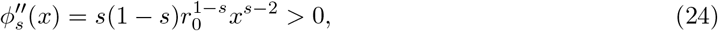

so *ϕ*_*s*_ is strictly convex for positive rates. Jensen’s inequality gives 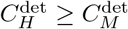, and hence

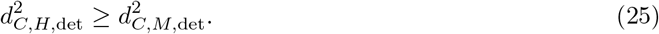

Again, equality holds only if 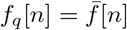 for every *q* and *n*. Thus a genuinely heterogeneous population gives strict detection improvement when *r*_0_ *>* 0.

The total firing-rate cost is the same for the heterogeneous and mean-copy populations,

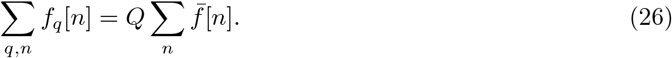

Consequently, with *L*^1^ metabolic cost, a heterogeneous population has better detection performance than a mean-copy population with the same metabolic cost under both independent Gaussian noise and independent Poisson noise with positive baseline.

### Existence of a homogeneous population preserving the information curve

The preceding results compare a heterogeneous population with *Q* copies of its arithmetic mean tuning curve. A different question is whether an improved Gaussian information curve *requires* a heterogeneous implementation. In the replicated Gaussian setting, the answer is no. One can construct a single homogeneous tuning curve whose shifted copies have exactly the same sampled Gaussian information tuning curve as the heterogeneous family.

We work on a circular grid *n* = 0, …, *K* − 1, with all indices taken modulo *K*. Let *f*_*q*_[*n*], *q* = 1, …, *Q*, denote the *Q* tuning-curve types in the heterogeneous family. We use the unitary discrete Fourier convention

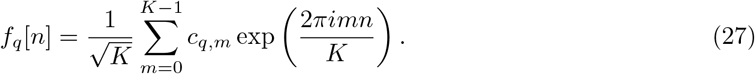

With this convention, Parseval’s identity gives

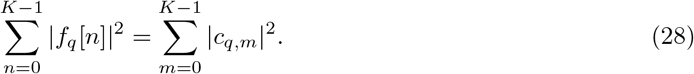

For independent Gaussian noise with variance *σ*^2^, the sampled information curve of the heterogeneous replicated population is

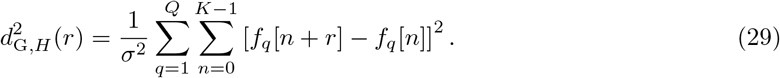

Using the Fourier expansion, this becomes

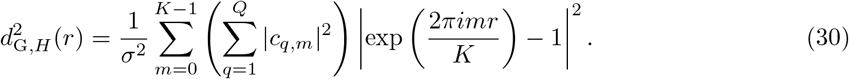

The zero-frequency term does not contribute. Thus the information curve depends only on the nonzero Fourier powers

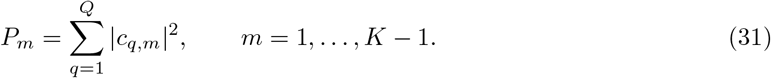

We now construct a homogeneous prototype *g*[*n*] with the same nonzero Fourier power. Let

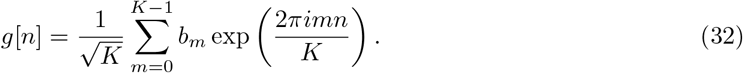

We require

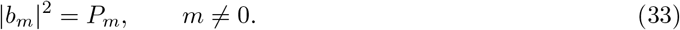

This condition is enough to guarantee that shifted copies of *g* have the same Gaussian information curve as the heterogeneous family.

The construction can be made explicit, real-valued, and even-symmetric. Since the original tuning curves are real, their Fourier coefficients satisfy

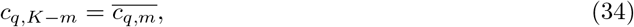

and therefore *P*_*K*−*m*_ = *P*_*m*_. Choose the nonzero Fourier coefficients of *g* to have zero phase:

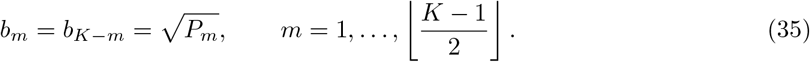

If *K* is even, the Nyquist coefficient is self-conjugate, and we choose

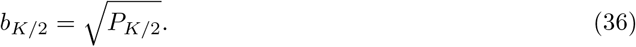

The zero-frequency coefficient *b*_0_ is not constrained by the information curve.

With these choices, the oscillatory part of the homogeneous prototype is real and even. If *K* is odd, define

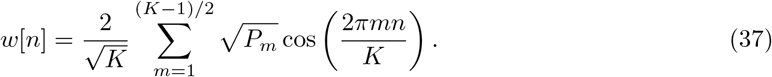

If *K* is even, define

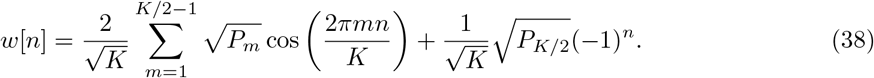

Then set

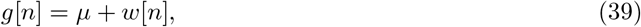

where 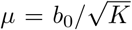 is a constant offset. By construction, *g*[− *n*] = *g*[*n*], so the prototype is even and centered at *n* = 0.

To make *g* a nonnegative firing-rate tuning curve, choose

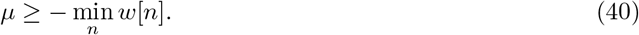

The smallest such choice makes the minimum response equal to zero. Adding any further constant offset leaves the information curve unchanged, because constants cancel in stimulus-stimulus differences.

The homogeneous population is obtained by placing shifted copies of this single prototype at the same preferred orientations:

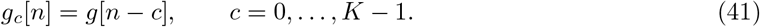

Its Gaussian information curve is

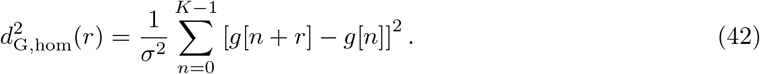

Using the Fourier coefficients *b*_*m*_, this becomes

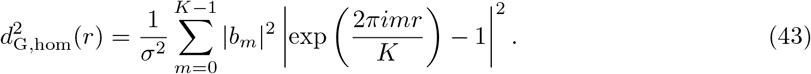

The zero-frequency term again vanishes. Since |*b*_*m*_|^2^ = *P*_*m*_ for all *m*≠0, Eqs. 30 and 43 imply

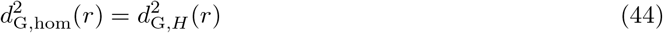

for every sampled separation *r*.

Thus a real, even-symmetric homogeneous prototype can be constructed explicitly by taking the square root of the summed nonzero Fourier power of the heterogeneous family and assigning zero Fourier phases. This is the finite-grid version of the Fourier-amplitude identity for homogeneous Gaussian populations [15]. An example of a construction is shown in Figure. 2. A similar construction can be derived for the Poisson noise case (not shown).

**Figure 1:**
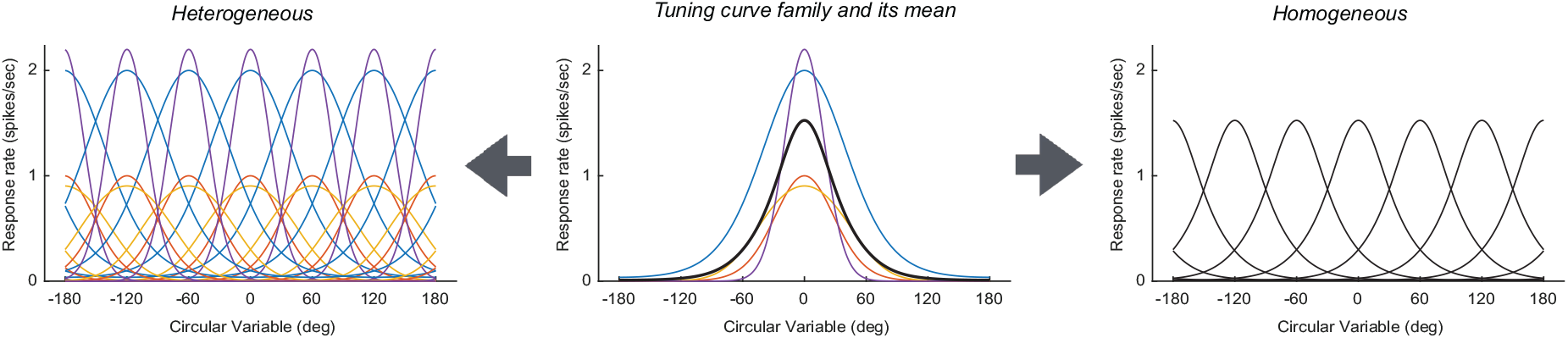
Schematic of the population construction. The center panel shows a family of *Q* tuning-curve types, *f*_*q*_(*θ*) in different colors, together with their mean tuning curve, 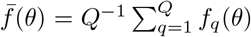, shown in black. The heterogeneous population on the left is formed by replicating every member of the family at each of *K* equally spaced preferred angles. The homogeneous comparison population on the right is formed by placing *Q* identical copies of the mean tuning curve at each preferred angle. Because these copies are identical, they are superimposed and appear as a single black curve at each location. Both populations therefore contain *QK* neurons and have the same total expected response, and hence the same *L*^1^ spike budget.

**Figure 2:**
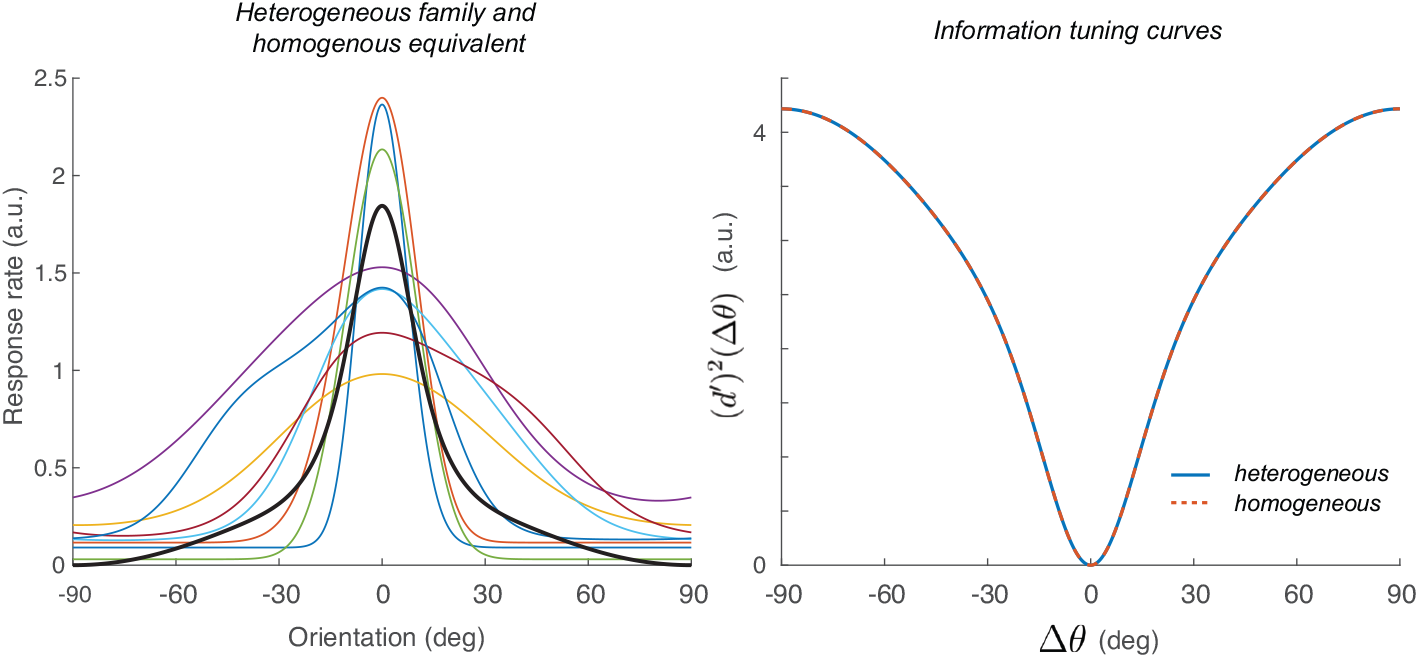
Example of a homogeneous population with the same Gaussian information tuning curve as a heterogeneous population. **Left:** Colored curves show a heterogeneous family of tuning curves centered at the same preferred angle. The black curve is the homogeneous prototype constructed by matching its nonzero Fourier power to the average Fourier power of the heterogeneous family. The corresponding homogeneous population contains the same number of neurons, implemented as identical copies of this prototype at each preferred angle. **Right:** Squared population distance as a function of angular separation for the heterogeneous and homogeneous populations. The curves overlap, verifying that Fourier-power matching produces identical information tuning curves at all sampled angular separations. This construction matches discrimination performance but does not necessarily preserve total *L*^1^ firing-rate cost.

It is difficult to determine the minimum metabolic cost required by the homogeneous population. The nonzero Fourier amplitudes are fixed by the information curve, but the Fourier phases and the DC component are not. Positivity of the tuning curves constrains the minimum allowable DC offset, and the resulting *L*^1^ cost depends on that number, which it is not trivial to compute and does not have a closed-form solution. This exercise shows that a given information curve can be implemented by a homogeneous population, while the prior results show that heterogeneity improves performance relative to copies of the mean tuning curve at fixed *L*^1^ cost.

### Average information constraint

#### Gaussian noise

The homogeneous Gaussian case also provides a useful way to understand how different norms of population activity determine average of the information tuning curve – that is, computing the average performance for different angular separations. We work on the standard circle 0 ≤ *θ <* 2*π*.

Let *f* be a nonnegative periodic tuning curve and define the Gaussian information tuning curve

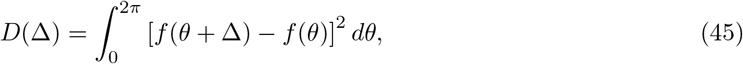

where angles are understood modulo 2*π*. Expanding the square gives

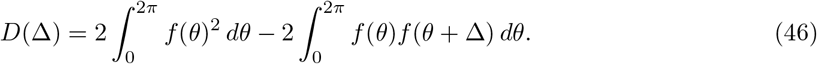

Thus *D*(Δ) is determined by the circular autocorrelation of the tuning curve.

Averaging over all angular separations,

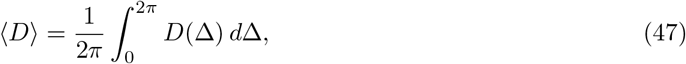

and using

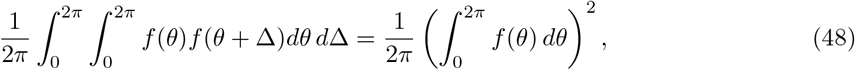

we obtain

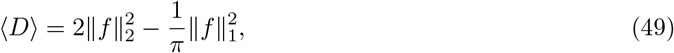

where

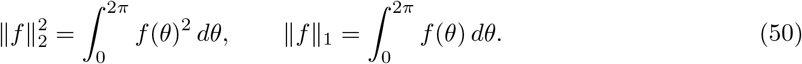

The second equality uses the nonnegativity of firing rates. Equivalently, defining the mean response

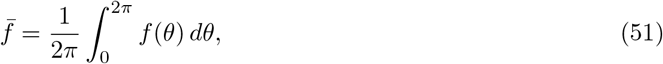

we can rewrite Eq. 49 as

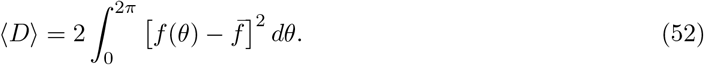

This is particularly informative as it show that the average information tuning curve is twice the *L*^2^ modulation energy of the tuning curve around its mean.

This identity clarifies the difference between *L*^2^ and *L*^1^ constraints. Under *L*^2^ normalization, 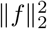 is fixed, and the average information depends only on the mean response. For example, if 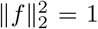 and

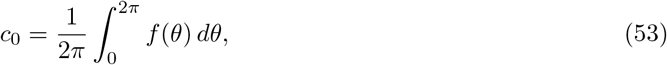

then Eq. 49 becomes

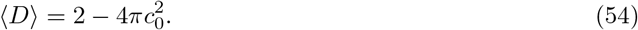

This is the average-performance constraint derived for homogeneous normalized populations in [15]. In this case, tuning curves with the same mean but different shapes have the same average information tuning curve.

Under an *L*^1^ spike-budget constraint, by contrast, the mean response is fixed:

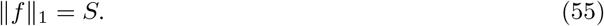

Equation 49 then becomes

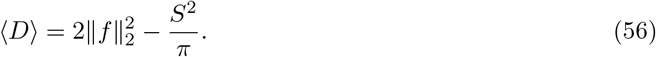

Thus *L*^1^ normalization does not impose a constant average information curve. Instead, at fixed total firing, average discrimination increases with the *L*^2^ energy of the tuning curve, or equivalently with its modulation around the mean. A flat tuning curve has zero average discrimination, whereas a more concentrated or strongly modulated tuning curve has larger average discrimination. In the absence of additional constraints, such as bounded peak rate, smoothness, bandwidth, or finite dimensionality, the *L*^1^-normalized continuous problem therefore has no analogous conservation law.

The same calculation has a finite-grid form. Let *f* [*n*], *n* = 0, …, *K* − 1, be a nonnegative tuning curve sampled on a circular grid, and define

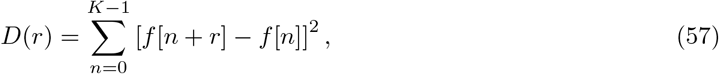

with indices taken modulo *K*. Then

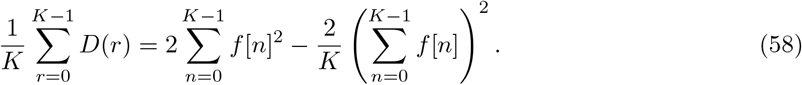

If the spike budget is fixed, 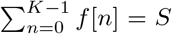, then the average information is again controlled by the squared modulation energy:

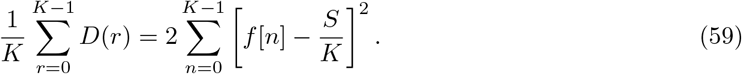

For a replicated heterogeneous family *f*_*q*_, the same identity applies term by term. If

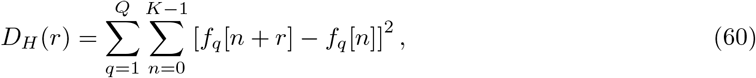

then

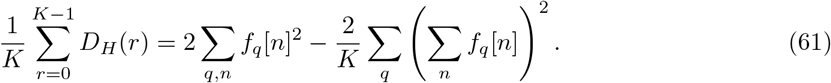

Writing

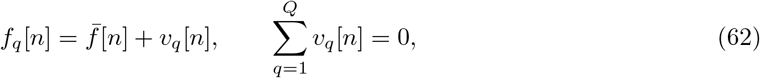

the average discrimination advantage relative to *Q* copies of the mean curve is

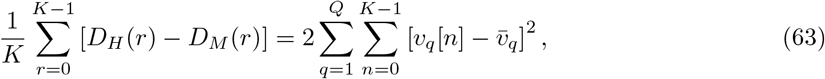

where

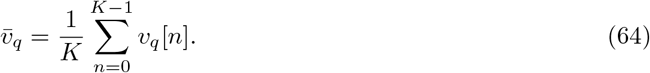

Thus the average Gaussian discrimination advantage of diversity is the non-DC modulation energy of the deviations from the mean. The *L*^1^-based diversity advantage and the *L*^2^-normalized average-information constraint are therefore two sides of the same autocorrelation calculation: *L*^2^ normalization fixes total power, whereas *L*^1^ normalization fixes the mean response and leaves modulation power available to improve average discrimination.

### Average Poisson information under an *L*^1^ constraint

It is possible to extend some of these results to the case of independent Poisson responses. Here, the finite-separation analogue of the Gaussian information curve should be defined from likelihoods. We consider a homogeneous tuning curve *f* (*θ*) ≥ 0 on the standard circle 0 ≤ *θ <* 2*π*. We also assume that the curve is even-symmetric about its peak, so after choosing coordinates with the peak at *θ* = 0,

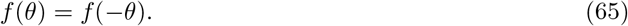

This requirement allows a major simplification in the Chernoff calculation. Let

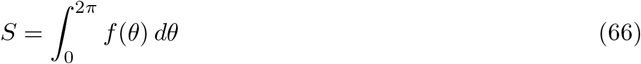

denote the total expected spike count. Two stimuli separated by Δ evoke rate functions *f* (*θ*) and *f* (*θ* + Δ). For a fixed Chernoff parameter *s* ∈ [0, 1], the Poisson Chernoff exponent is

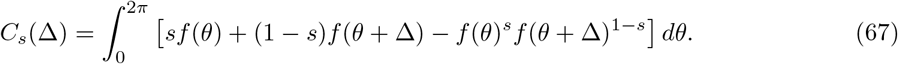

Because circular shifts preserve the integral of *f*, this simplifies to

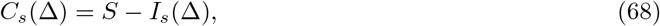

where

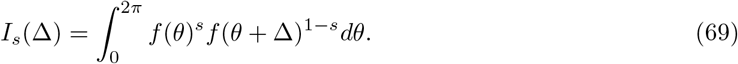

The exact Chernoff information is

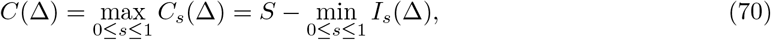

and the Chernoff-equivalent finite-separation measure is

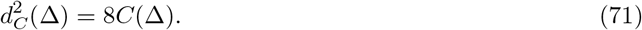

The even-symmetry assumption makes the Chernoff optimization explicit. First, by the change of variables *u* = −*θ* − Δ, and using *f* (*u*) = *f* (−*u*), we obtain

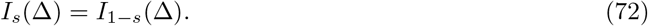

Second, *I*_*s*_(Δ) is log-convex as a function of *s*, by Hölder’s inequality. Therefore, since *I*_*s*_(Δ) is symmetric around *s* = 1*/*2, its minimum occurs at *s* = 1*/*2. Consequently,

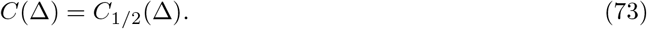

Thus, for even-symmetric tuning curves, the exact Poisson Chernoff information is the Bhattacharyya distance:

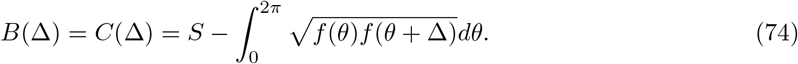

Equivalently,

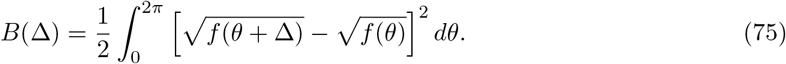

This form emphasizes that, in the symmetric Poisson case, finite-separation information is a squared Euclidean distance in square-root-rate coordinates.

Averaging over all angular separations gives

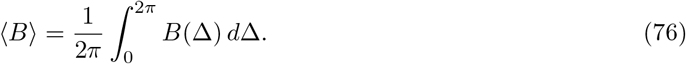

Using

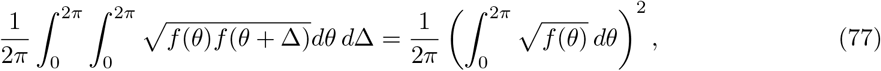

we obtain

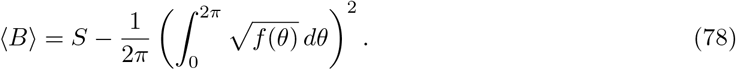

Therefore the average Chernoff-equivalent *d*^′2^ is

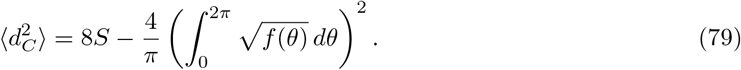

This is the closest Poisson analogue of the Gaussian average-information identity. Under a fixed *L*^1^ spike budget, *S* is fixed, but the square-root moment

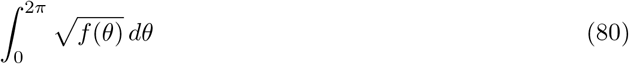

is not fixed. Thus average Poisson information is not conserved under *L*^1^ normalization. A flat tuning curve minimizes the average information. Indeed, if *f* (*θ*) = *S/*(2*π*), then

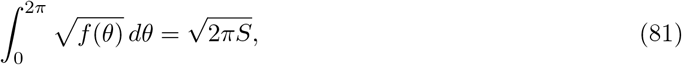

and Eq. 78 gives ⟨*B*⟩ = 0. As the tuning curve becomes more concentrated at fixed total spike count, the square-root moment decreases and the average information increases. In the limit of increasingly sparse responses at fixed *S*, ⟨*B*⟩ approaches *S*, so

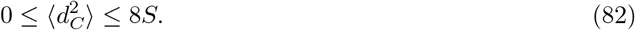

## Discussion

We considered a simplified model of coding a circular variable that has several important limitations. First, heterogeneity is introduced in a highly structured manner that permits analytical treatment but is not intended as a realistic model of cortical tuning diversity. Second, we use total expected spike count as a tractable proxy for metabolic cost. Cortical energy expenditure also includes postsynaptic currents, maintenance of ionic gradients and resting potentials, transmitter release, and transmitter recycling, and current energy budgets indicate that action potentials constitute only one—and not necessarily the largest—component of signaling-related energy expenditure [16–18]. Within these limitations, however, the model allows us to isolate how tuning diversity affects coding performance under a fixed *L*^1^ spike-count constraint. Our main comparison is between a replicated heterogeneous family of tuning curves and an equally sized population composed of replicated copies of its arithmetic mean tuning curve. The two populations have identical total expected responses and therefore the same value of our metabolic-cost proxy. Any difference in performance consequently reflects the structure of the tuning curves rather than a difference in spike budget.

For discrimination peformance we obtained similar results for Gaussian and Poisson noise models. In the Gaussian case, raw *d*^′2^ is a squared Euclidean distance between population response vectors. Here, decomposing each tuning curve into the mean curve plus a mean-preserving deviation shows that the heterogeneous information tuning curve equals the mean-copy information tuning curve plus a nonnegative squared-difference term. Equality occurs only when heterogeneity is achieved by DC shifting the tuning curves. Other than this peculiar condition all other cases lead to strict gains at least at some angular separations. For Poisson noise we measure discrimination by the Chernoff information. We obrain a similar gain, but the equality conditions are different. Pure multiplicative gain diversity, *f*_*q*_ = *a*_*q*_*g*, is neutral for the Poisson Chernoff comparison with copies of the mean, whereas shape diversity is beneficial whenever it changes shape.

For detection performance we measured how the population response differed from that of a uniform baseline. Detection fixes an origin in response space: the no-stimulus response with baseline rate *r*_0_. For Gaussian detection, the comparison with the mean-copy population is governed by the squared distance from this baseline, so any genuine heterogeneity on the sampled grid gives a strict improvement. For Poisson detection, strict convexity of the one-variable Chernoff exponent gives the same conclusion: equality with the mean-copy population occurs only when all tuning curves coincide with the mean on the sampled grid.

Although diversity improves performance relative to copies of the mean tuning curve, this does not imply that the resulting Gaussian information tuning curve *requires* a heterogeneous implementation. Indeed, we showed how a homogeneous population can be constructed given a desired information tuning curve by matching that Fourier power. This is a finite-grid version of the Fourier-amplitude identity for homogeneous Gaussian populations described earlier [15].

Finally, the relationship between population norms and the average of the information tuning curve provides additional insights. For Gaussian noise, we obtained

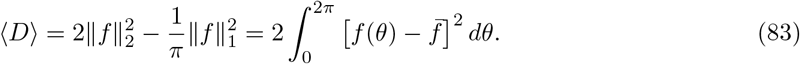

Thus *L*^2^ normalization fixes total power and makes the average information depend only on the mean response, whereas *L*^1^ normalization fixes the mean response and leaves the *L*^2^ modulation energy free to vary. On the finite grid, the average Gaussian discrimination gain of the heterogeneous family over the mean-copy population is exactly twice the non-DC *L*^2^ energy of the deviations 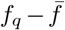. The Poisson noise analogue of this result has the same form after transforming to square-root-rate coordinates in the symmetric case. If *f* is even-symmetric, so that the Chernoff optimum occurs at *s* = 1*/*2, then we obtained

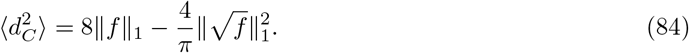

The central message of these calculations is that exact homogeneity is not an evolutionary ‘attractor’ within the coding problem considered here. At a fixed spike budget, perturbations of a homogeneous tuning curve that preserve the mean improve discrimination or detection performance, except in restricted equality cases. If establishing identical tuning curves also requires greater developmental coordination or wiring precision than allowing variability, then both coding performance and construction cost should bias selection toward heterogeneous populations. This possibility is consistent with large-scale, anatomically constrained models of V1, in which the distribution of many response properties—including orientation selectivity, simple and complex cell responses, and direction selectivity—emerge from shared cortical circuitry acting on thalamic inputs [19–22]. From this generative perspective, receptive-field diversity may be the inexpensive developmental default, whereas producing and maintaining a homogeneous population would require additional control. If such readily generated diversity is also beneficial, or at least not detrimental, to coding, evolution need not design heterogeneity explicitly; it need only avoid suppressing it. Tuning diversity may therefore reflect both efficient coding and the economical use of the circuit-building mechanisms available to cortex.

It may be possible to test these ideas in more complex evolutionary simulations that relax the assumptions we imposed to make the problem mathematically tractable. Recent work has demonstrated that sensory morphology, neural processing, and behavior can be co-evolved in embodied agents, with different task demands producing distinct visual solutions [23]. A similar evolutionary framework could be extended to treat tuning curves and cortical circuit costs as heritable traits, define fitness in terms of discrimination, detection, metabolic expenditure, and wiring cost, and ask whether diversity emerges spontaneously, under what conditions homogeneity persists, and how environmental demands shape the resulting distribution of tuning properties.

## Acknowledgments

This work was inspired by the Ph.D. dissertation of Sonica Saraf at New York University. I thank Bob Shapley for comments on an earlier version of this manuscript. This was was supported by National Institutes of Health (NIH) Grants EY034488, NS116471, and EY036219 to D.L.R. and EY023871 to Joshua Trachtenberg and D.L.R.

